# Role of the high-affinity leukotriene B_4_ receptor signaling in fibrosis after unilateral ureteral obstruction in mice

**DOI:** 10.1101/390914

**Authors:** Mariko Kamata, Hideki Amano, Yoshiya Ito, Tomoe Fujita, Kanako Hosono, Kouju Kamata, Yasuo Takeuchi, Takehiko Yokomizo, Takao Shimizu, Masataka Majima

**Affiliations:** Departments of Pharmacology, Kanagawa 252-0374, Japan; Departments of Nephrology Kitasato University School of Medicine, Kanagawa 252-0374, Japan; Department of Molecular Pharmacology, Kitasato University Graduate School of Medical Sciences, Kanagawa 252-0374, Japan; Department of Pharmacology and Toxicology, Dokkyo Medical University School of Medicine, Tochigi 321-0293, Japan; Sagamiono Medical and Kidney Clinic, Kanagawa 252-0303, Japan; Department of Biochemistry, Juntendo University School of Medicine, Tokyo 113-8421, Japan; Department of Lipidomics, Faculty of Medicine, University of Tokyo, Tokyo, 153-8902, Japan; Department of Lipid Signaling, National Center for Global Health and Medicine, Tokyo 162-0052, Japan

**Keywords:** BLT1, Leukotriene B_4_, Fibrosis, Unilateral ureteral obstruction, Macrophage, Fibroblast, TGF-β

## Abstract

Leukotriene B_4_ (LTB_4_) is a lipid mediator that acts as a potent chemoattractant for inflammatory leukocytes. Kidney fibrosis is caused by migrating inflammatory cells and kidney-resident cells. Here, we examined the role of the high-affinity LTB_4_ receptor BLT1 during development of kidney fibrosis in wild-type (WT) mice and BLT1 knockout (BLT1^-/-^) mice with unilateral ureteral obstruction (UUO). We found elevated expression of 5-lipoxygenase (5-LOX), which generates LTB_4_, in the renal tubules of WT and BLT1^-/-^ UUO mice. Accumulation of immunoreactive type I collagen in UUO kidneys of WT mice increased over time; however, the increase was less prominent in BLT1^-/-^ mice. Accumulation of S100A4-positive fibroblasts also increased temporally in WT UUO kidneys, but was again less pronounced in those of BLT1^-/-^ mice. The same was true of mRNA encoding transforming growth factor-β (TGF)-β and fibroblast growth factor (FGF)-2. Finally, accumulation of F4/80-positive macrophages, which secrete TGF-β, also increased temporally in WT UUO and BLT1^-/-^ kidneys, but to a lesser extent in the latter. Following LTB_4_ stimulation *in vitro*, macrophages showed increased expression of mRNA encoding TGF-β/FGF-2 and Col1a1, whereas L929 fibroblasts showed increased expression of mRNA encoding α smooth muscle actin (SMA). Bone marrow (BM) transplantation studies revealed that the area positive for type I collagen was significantly smaller in BLT1^-/-^-BM→WT UUO kidneys than in WT-BM→WT kidneys. Thus, LTB_4_-BLT1 signaling plays a critical role in fibrosis in UUO kidneys by increasing accumulation of macrophages and fibroblasts. Therefore, blocking BLT1 may prevent renal fibrosis.

## Introduction

Unilateral ureteral obstruction (UUO) is an experimental animal model of renal fibrosis that mimics the pathogenesis of chronic obstructive nephropathy in humans. The hydrostatic pressure resulting from the obstruction triggers expression of chemokines in tubular epithelial cells [1], followed by increased interstitial capillary permeability [2], infiltration by interstitial inflammatory cells [3], myofibroblast activation, and extracellular matrix deposition [4–6]. Progressive fibrosis, loss of renal parenchyma due to capillary rarefaction [4], and tubular cell death via apoptosis and necrosis [7] also occur. Despite these severe changes in the obstructed kidney, the animal remains healthy because the contralateral kidney is fully functional. Indeed, unlike renal ablation models [8], UUO model mice do not have uremia. Therefore, the UUO model is ideal for studying the histopathological and molecular changes underlying tubulointerstitial damage, a process that closely resembles deterioration of renal function in humans with chronic kidney disease [9–11].

Macrophages are rarely present in the healthy renal cortex [12, 13]. However, within hours of ureteral obstruction, a large number of blood-derived macrophages accumulate in the tubulointerstitial space [14]. This cellular infiltration is preceded by local expression of chemokines [1, 15], chemokine receptors [1], and adhesion molecules [16, 17]. Despite the accumulation of strong correlative data, there are few functional studies describing the role of these infiltrating macrophages and lipid mediators in UUO-induced fibrosis. A previous report shows that prostaglandin (PG) E_2_, a major metabolite of arachidonic acid, suppresses tubulointerstitial fibrosis via EP4 [18]. Furthermore, EP4 signaling suppresses accumulation of macrophages in the kidneys following induction of UUO. A recent report suggests that bone marrow (BM)-derived macrophages, which express c phospholipase (PLA)_2_α, upstream of the 5-lipoxygenase (5-LOX) pathway, exacerbate fibrosis in the UUO kidney [19].

Leukotrienes (LTs) are metabolites of arachidonic acid that are generated via the 5-LOX (EC 1.13.11.34, 5-LOX) pathway. LTB_4_ is a well-characterized and potent chemoattractant for leukocytes, particularly neutrophils and monocytes [20]; as such, it plays a pivotal role in the pathogenesis of inflammatory and immune diseases such as asthma [21], sepsis [22], and atherosclerosis [23, 24]. Previously, we showed that LTB_4_ is a potent inducer of neutrophil extravasation into the interstitial space in certain *in vivo* models [25–28]. LTB_4_ exerts its biological activity through two distinct receptors: LTB_4_ receptor type-1 (BLT1), a high-affinity LTB_4_ receptor highly expressed in leukocytes, and BLT2, a low-affinity LTB_4_ receptor expressed more ubiquitously than BLT1 in human tissues [29–31].

Hemodynamic changes, which are dependent on the 5-LOX pathway, were described in a rat model of bilateral ureteral obstruction [32]. Although, blocking LTB_4_ activity reduces fibrosis in bleomycin-treated lungs [33], the role of BLT1 signaling in UUO-induced fibrosis remains unclear.

Here, we examined the role(s) of BLT1 signaling in development of fibrosis in a BLT1 knockout (BLT1^-/-^) mouse model of UUO [34]. We noted significantly less accumulation of type I collagen in kidneys of BLT1^-/-^ mice with UUO than in those of wild-type (WT) mice. We concluded that LTB_4_-BLT1 signaling plays a role in tubulointerstitial fibrosis of the kidney, possibly via upregulation of TGF-β and increased recruitment of myofibroblasts and fibroblasts. Thus, blocking BLT1 signaling may prevent fibrosis in those with chronic kidney disease.

## Materials and methods

### Animals and Surgery

BLT1^-/-^ mice were developed as described previously [34]. Male C57BL/6 WT mice and BLT1^-/-^ mice (8 weeks old) were used. UUO surgery was performed under inhalation anesthesia of isoflurane mixed with air and its adequacy was monitored from the disappearance of the pedal withdrawal response. A median abdominal incision was made, and the left proximal ureter was ligated at two points using 3-0 silk. The incision was closed with wound clips (AUTOCLIP, 9 mm; ALZET, Cupertino, CA, USA). Sham-operated mice had the ureter exposed but not ligated [36]. All experiments were performed in accordance with the guidelines for animal experiments established by the Kitasato University School of Medicine (2018-166) and conformed to the Guide for the Care and Use of Laboratory Animals published by the US National Institutes of Health (NIH Publication No. 85–23, revised 1996). The mice were maintained at constant humidity (60 ± 5 %) and temperature (22 °C ± 1) on a 12-h light/ dark cycle. All animals were provided with food and water ad libitum. The total number of mice used in this experiment is 166. The number of mice per group is from 4 to 20. At the end point of the experiments, mice were sacrificed under inhalation anesthesia of isoflurane mixed with air. Mice exhibiting symptoms of infection including suppressed appetite, purulent discharge from the wound were removed from the study prior to the study endpoint.

### Tissue harvesting

Kidney samples were collected on Days 0, 1, 3, 5, 7, 10, and 14 after UUO. Day 0 kidney samples were collected without the need for surgical procedures. All mice were anesthetized with isoflurane and perfused with PBS via the left ventricle. The left kidney was harvested immediately and cut into transverse sections for RT-PCR, paraffin embedding, freezing, and Sircol collagen assays.

### Histological examination

Kidney tissues were fixed overnight at 4°C in 4% paraformaldehyde and embedded in paraffin. Paraffin-embedded tissues were cut into 4 μm sections and stained with H&E and Sirius red. Kidney cortex thickness was measured by investigators blinded to treatment arm using Image J software using.

### Immunofluorescence staining

Unfixed kidney tissues were frozen immediately in liquid nitrogen. Samples were cut into 4 μm sections from the cortical side, blocked with 1% BSA in PBST (0.1% Triton X-100 in PBS) for 1 h at room temperature, and incubated overnight at 4°C with an anti-type I collagen antibody (1:100 dilution; Abcam, Cambridge, UK; ab21286). After washing in PBS, the sections were incubated for 1 h at room temperature with Alexa Fluor^®^ 488-Donkey anti-rabbit IgG (1:500 dilution; Molecular Probes, Eugene, OR, USA). Five randomly selected cortical interstitial fields from each animal were photographed (at ×400 magnification), excluding the glomeruli and large vessels. The immunoreactive interstitial area was calculated using Image J software and expressed as a percentage of the total area. Periodate-lysine-paraformaldehyde tissues were fixed for 2 h at 4°C, frozen in liquid nitrogen, cut into 10 μm sections, and stained as described above with one of the following primary antibodies: anti-5-LOX (1:100 dilution; Novus Biologicals, Littleton, CO, USA; NB 100-92138), anti-CXCL12 (1:100 dilution; eBioscience, San Diego, CA, USA; 14-7992), or anti-F4/80 (1:200 dilution; Santa Cruz Biotechnology, Inc., Dallas, TX, USA; sc-52664). After washing in PBS, the sections were incubated for 1 h at room temperature with one of the following secondary antibodies: Alexa Fluor^®^ 488-Donkey anti-Rabbit IgG (1:500 dilution, Molecular Probes) or Alexa Fluor^®^ 568-Donkey anti-Rat IgG (1:500 dilution, Molecular Probes).

### Immunohistochemistry

Paraffin-embedded tissues were cut into 4 μm sections, deparaffinized, and rehydrated. Endogenous peroxidase was quenched by immersion for 30 min in a 1% solution of hydrogen peroxide in methanol. After washing in ion-exchanged water, antigen retrieval was performed by microwaving three times for 5 min in citrate buffer solution (pH 6.0). The sections were then incubated for 10 min with Protein Block, Serum-Free (DAKO, Glostrup, Denmark), followed by an overnight incubation at 4°C with an anti-S100A4 antibody (1:400 dilution; Abcam, ab27957). After washing in PBS, the sections were incubated for 30 min at room temperature with N-Histofine^®^ Simple Stain^™^ MAX PO (R) (Nichirei Biosciences, Inc., Tokyo, Japan). Immune complexes were then detected with 3, 3’-diaminobenzidine tetrahydrochloride (DAB), and sections were counterstained with methyl green. The number of S100A4-positive interstitial cells in five random cortical fields (×200 magnification) per sample was counted. All images were captured by a Biozero BZ-9000 series microscope (Keyence, Tokyo, Japan).

### Sircol collagen assay

The total amount of soluble collagen was measured using a Sircol collagen assay kit (Biocolor, Antrim, UK). Kidney samples were frozen immediately with in liquid nitrogen immediately and stored at −80°C until use. All measurements were performed in duplicate and results were expressed as μg of collagen/mg of kidney cortex.

### Real-time RT-PCR

Total RNA was extracted from decapsulated kidney tissues using TRIzol^®^ reagent (Gibco-BRL; Life Technologies, Rockville, MD, USA), and single-stranded cDNA was generated from 1 μg of total RNA via reverse transcription using the ReverTra Ace^®^ qPCR RT Kit (TOYOBO CO., LTD., Osaka, Japan), according to the manufacturer’s instructions. Real-time PCR was performed using SYBR^®^ Premix Ex Taq™ II (Tli RNaseH Plus; Takara Bio, Inc., Shiga, Japan). The gene-specific sequences are described in Table 1. Expression of target genes was normalized to that of GAPDH.

**Table 1.**
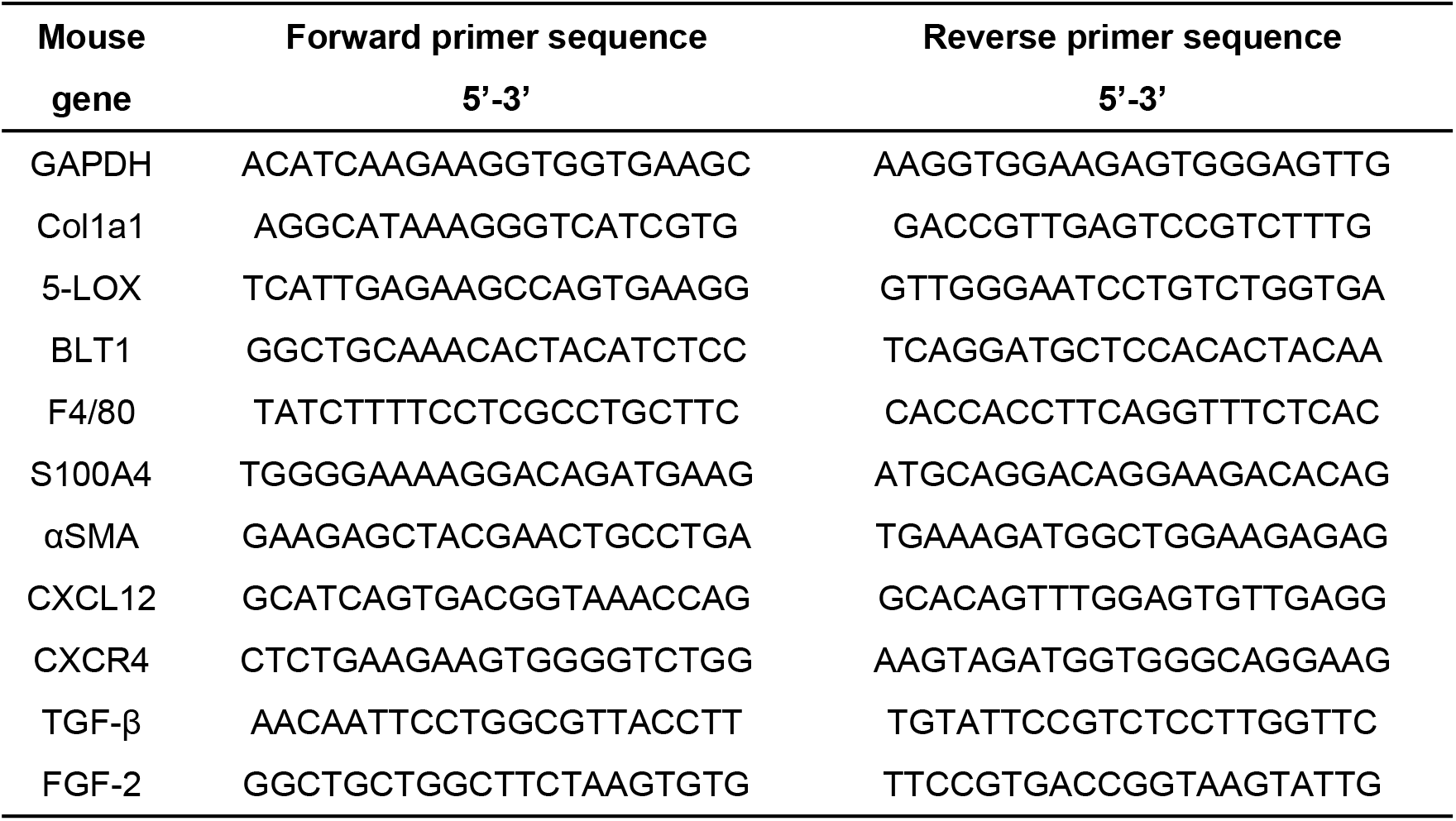
Primers used for reverse transcription and quantitative PCR.

### Collection of peritoneal macrophages

Thioglycolate-induced peritoneal macrophages were collected from 8–12-week-old C57/BL6 WT mice. In brief, 2 ml of 4% thioglycolate medium was injected into the peritoneal cavity. After 3 days, the peritoneal cavity was washed three times with 5 ml of PBS. Cells in the lavage fluid were washed and suspended in RPMI 1640 medium containing 10% FCS and then placed in 12-well culture plates (1 × 10^6^ cells/well). After incubation at 37°C in 5% humidified CO_2_ for 16 h, the plates were washed with PBS to remove non-adherent cells. Approximately 60% of cells remained adherent and were used for subsequent experiments.

### Cell culture and treatments

At 16 h after plating, cells were washed twice with PBS and incubated for 2 h in serum-free medium. Cells were then stimulated for 12 h with LTB_4_ (0.1, 1, or 10 nM) or serum-free medium (control). Total mRNA was isolated from cells using TRIzol^®^ reagent, and mRNA expression was measured by real-time RT-PCR.

Thioglycolate-induced peritoneal macrophages (after removal of non-adherent cells) were stimulated with LTB_4_, and expression of TGF-β, FGF-2, αSMA, and Col1a1 mRNA was measured. Murine fibroblasts (L929) were purchased from the Cell Bank at RIKEN BioResource Center (Ibaraki, Japan). Cells were suspended in DMEM containing 10% FCS, plated in 6-well culture plates (3 × 10^5^ cells/well), and stimulated with LTB_4_ and TGF-β, and expression of TGF-β, FGF-2, αSMA, and Col1a1 mRNA was measured.

### BM transplantation

BM transplantation experiments were carried out as described previously [37]. In brief, donor BM was obtained by flushing the femoral and tibial cavities of WT mice and BLT1^-/-^ transgenic mice with PBS. The flushed BM cells were dispersed and resuspended in PBS at a density of 1 × 10^6^ cells/100 μl. Both WT and BLT1^-/-^ mice were lethally irradiated with 9.5 Gy X-rays using an MBR-1505 R X-ray irradiator (Hitachi Medico, Tokyo, Japan) equipped with a filter (copper, 0.5 mm; aluminum, 2 mm). The cumulative radiation dose was monitored. BM mononuclear cells from WT and BLT1^-/-^ mice (2 × 10^6^ cells/200 μl) were transplanted into irradiated WT and BLT1^-/-^ mice via the tail vein.

### Statistical analysis

All results are expressed as the mean ± SEM. Comparisons between two groups were performed using Student’s *t* test. Comparisons between multiple groups were performed using one-way ANOVA, followed by Tukey’s *post-hoc* test. P values <0.05 were considered statistically significant.

## Results

### Development of fibrosis in WT mouse kidneys after UUO

Following induction of UUO, the thickness of the kidney cortex in WT mice decreased gradually (Fig. 1A) and was significantly smaller than that in sham-operated mice at Days 7 and 14 (Day7: 1.14±0.05 vs.1.62±0.04 mm, P<0.0001, Day14: 0.68±0.05 vs.1.59±0.03 mm, P<0.0001, Fig. 1B). Sirius red staining demonstrated that areas of collagen deposition around dilated renal tubules increased in a temporal manner (Fig. 1C), and the Sircol collagen assay showed that collagen levels on Days 7 and 14 were significantly higher than sham-operated mice (Day7: 8.85±1.20 vs.3.61±0.55μg/mg of kidney weight, P=0.018, Day14: 11.62±1.15 vs.3.47±0.16μg/mg, P= 0.0007, Fig. 1D).

**Figure 1.**
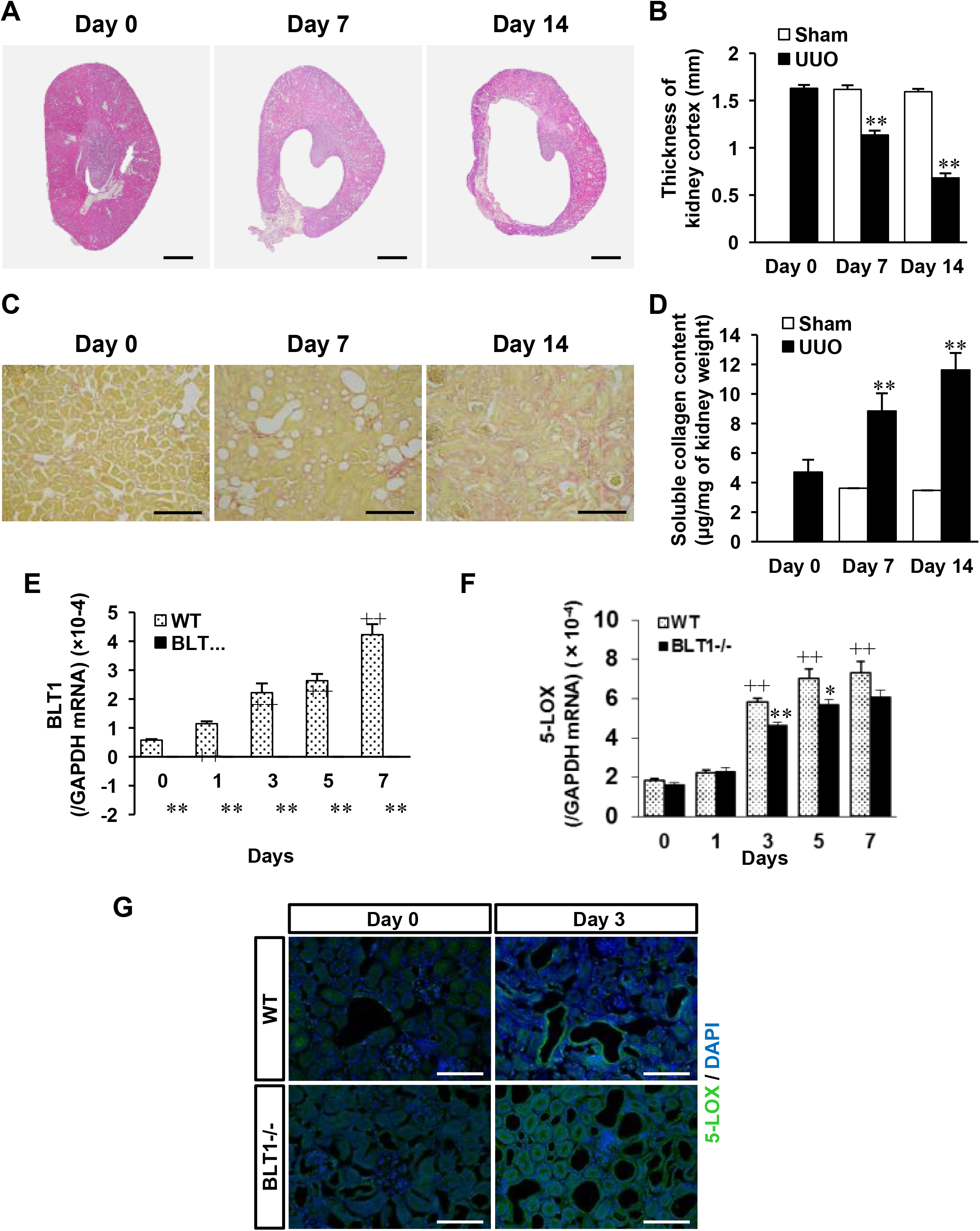
Effect of BLT1 and 5-LOX on development of fibrosis after UUO. A) Photomicrographs showing H&E-stained transverse sections of WT kidney. Scale bars = 500 μm. B) Changes in thickness of the WT UUO kidney cortex. Data are expressed as the mean ± SEM (n=4 mice/group). **P<0.01, *vs*. sham. C) Typical images showing Sirius red staining of WT kidney cortex after UUO. Scale bars = 200 μm. D) Soluble collagen in the WT UUO kidney cortex measured in the Sircol collagen assay. Data are expressed as the mean ± SEM (n=6 mice/group). **P<0.01, *vs*. Day 0. E,F) Expression of mRNA encoding BLT1 (E) and 5-LOX (F) in UUO kidneys, as measured by real-time RT-PCR. Data are expressed as the mean ± SEM (n=8-15 mice per group). *P<0.05 and **P<0.01, *vs*. WT; +P<0.05 and ^++^P<0.01, *vs*. Day 0 WT. G) Images of sections of WT and BLT1^-/-^ UUO kidney cortex immunostained with an anti-5-LOX antibody on Days 0 and 3. Epithelial cells in dilated renal tubules and some interstitial cells stained positive for 5-LOX. Scale bars = 200 μm.

### Expression of BLT1 and 5-LOX increases in UUO kidneys

To study the role of LTB_4_-BLT1 signaling in the UUO kidney, we examined expression of 5-LOX (an enzyme upstream of LTB_4_) and BLT1. Expression of BLT1 mRNA in WT UUO kidneys on Day 1 was markedly higher than that on Day 0 (P<0.0001), while BLT1 mRNA levels in BLT1^-/-^ mice were negligible throughout the experimental period (Fig. 1E). By contrast, expression of 5-LOX increased in both WT and BLT1^-/-^ mice from Day 3, although levels in BLT1^-/-^ mice were significantly lower than those in WT mice at Days 3 and 5 (Day3: P=0.0002, Day5: P=0.03, Fig. 1F). Immunostaining of WT and BLT1^-/-^ kidneys for 5-LOX at Day 3 after induction of UUO revealed that dilated tubule epithelial cells in WT mice were positive, as were some interstitial cells. By contrast, there were fewer 5-LOX-positive cells in dilated tubules and tubulointerstitial areas of BLT1^-/-^ kidneys (Fig. 1G).

### Tubulointerstitial fibrosis in BLT1^-/-^ mice is reduced after UUO

To examine the role of BLT1 signaling in collagen accumulation, we examined kidney fibrosis in WT and BLT1^-/-^ mice after UUO. Immunostaining of type I collagen increased in WT kidneys after induction of UUO (Fig. 2A). Quantitative analysis of the immunoreactive renal interstitial area revealed an increase in the percentage positive area after induction of UUO in both WT and BLT^-/-^; however, the area of type I collagen was significantly lower in BLT1^-/-^ mice than in WT mice from Day 3 (Day3: 5.95±0.05 vs.5.13±0.11 %, P<0.001, Day5: 7.36±0.30 vs.5.66±0.10, P=0.004, Day7: 12.9±0.99 vs.9.46±0.29, P=0.02, WT vs. BLT1, respectively; Fig. 2B). Furthermore, expression of mRNA encoding Col1a1 in WT and BLT^-/-^ UUO kidneys increased in a temporal manner, but was significantly lower in BLT1^-/-^ mice from Day 1 (Day1: <0.0001, Day3: P<0.0001, Day5: P=0.008, Day7: P=0.0002, WT vs. BLT1, respectively; Fig. 2C).

**Figure 2.**
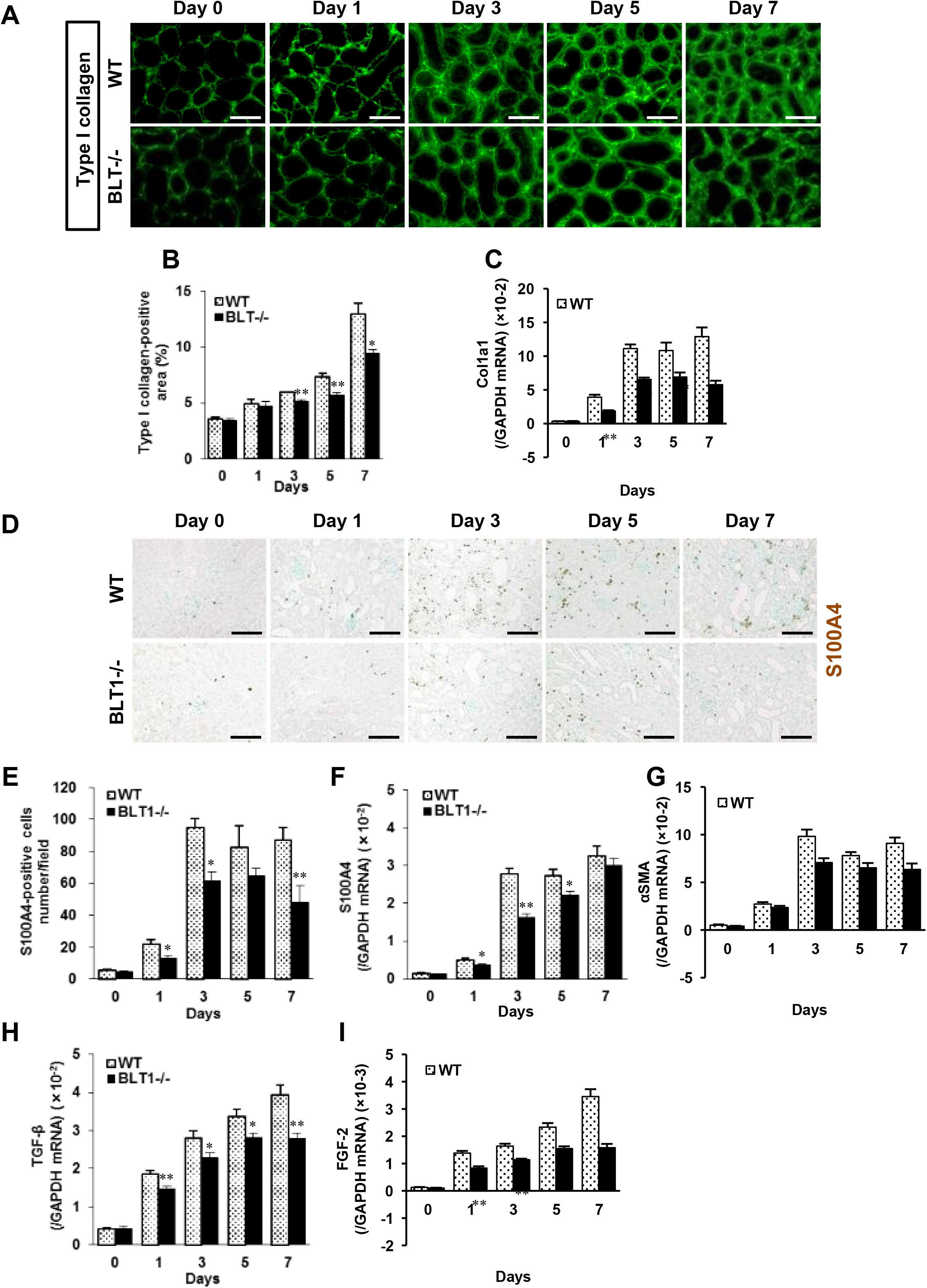
Tubulointerstitial fibrosis is less severe in BLT1^-/-^ mice following UUO. A) Representative images of kidney cortex from WT and BLT1^-/-^ mice immunostained with an anti-type I collagen antibody. Scale bars = 50 μm. B) Temporal changes in the area of immunoreactive collagen within tubulointerstitial spaces (expressed as % total area, excluding glomeruli and large vessels). Data are expressed as the mean ± SEM (n=4 mice/group). *P<0.05 and **P<0.01, *vs*. WT. Expression of Col1a1 mRNA (C) in WT and BLT1^-/-^ mice kidneys, as measured by real-time PCR. Data are expressed as the mean ± SEM (n=8-20 mice per group). *P<0.05 and **P<0.01, *vs*. WT. D) Representative images showing S100A4 staining in WT and BLT1^-/-^ kidneys after UUO. Scale bars = 100 μm. E) Changes in the number of S100A4-positive cells in WT and BLT1^-/-^ UUO kidneys. The number of S100A4-positive cells was significantly lower in BLT1^-/-^ UUO kidneys. Data are expressed as the mean ± SEM (n=4 mice/group). F, G) Expression of mRNA encoding S100A4 (F) and αSMA (G) in UUO kidneys, as measured by real-time PCR. Data are expressed as the mean ± SEM (n=8-18 mice per group). *P<0.05 and **P<0.01, *vs*. WT. H, I) Expression of mRNA encoding TGF-β (H) and FGF-2 (I) in mouse kidneys after induction of UUO. Data are expressed as the mean ± SEM (n=8-20 mice per group). *P<0.05 and **P<0.01, *vs*. WT mice.

### BLT1-dependent accumulation of fibroblasts in UUO kidneys

S100A4-positive fibroblasts accumulated in the interstitial tissues of WT and BLT1^-/-^ UUO kidneys (Fig. 2D). The number of S100A4-positive cells in immunohistochemical specimens from WT UUO kidneys increased in a time-dependent manner; however, there were significantly fewer S100A4-positive cells in BLT1^-/-^ UUO kidneys than in WT kidneys at Days 1, 3, and 7 (Day1: 21.9±2.9 vs.13.0±1.8 cells/field, P=0.04, Day3: 94.7±5.7 vs.61.8±5.4, P=0.006, Day7: 87.1±7.9 vs.48.1±10.8, P=0.03, WT vs. BLT1, respectively; Fig. 2E). In addition, expression of S100A4 mRNA in WT UUO kidneys increased, whereas that in BLT1^-/-^ UUO kidneys was suppressed, at Days 1, 3, and 5 (Day1: P=0.017, Day3: P<0.0001, Day5: P=0.02, WT vs. BLT1, respectively; Fig. 2F). Changes in S100A4 mRNA levels mirrored changes observed upon immunohistochemical analysis. Expression of mRNA encoding αSMA, a myofibroblast marker, increased in WT UUO kidneys; however, this increase was significantly lower in BLT1^-/-^ UUO than in WT kidneys after Day 3 (Day3: P=0.008, Day5: P=0.012, Day7: P=0.003, WT vs. BLT1, respectively; Fig. 2G).

### Upregulation of TGF-β and FGF-2 in UUO kidneys is BLT1-dependent

Several growth factors are expressed in UUO kidneys [10, 38]. Real-time PCR showed that expression of TGF-β mRNA in WT UUO kidneys increased from Day 1 post-induction, whereas the increases in BLT1^-/-^ mice were significantly lower (Day1: P=0.006, Day3: P= 0.03, Day5: P=0.02, Day7: P=0.002, WT vs. BLT1, respectively; Fig. 2H). The same was true for FGF-2 mRNA (Day1, 3, 5, 7: P<0.001, WT vs. BLT1, respectively; Fig. 2I).

### Accumulation of macrophages in UUO kidneys is BLT1-dependent

Interstitial macrophages promote fibrosis in the UUO kidney [39]. Therefore, we asked whether macrophages accumulate in the interstitial spaces within UUO kidneys. Immunofluorescence analysis showed that fewer F4/80-positive macrophages accumulated in interstitial tissues of BLT1^-/-^ UUO kidneys than in those of WT kidneys (Fig. 3A). The macrophage density in BLT1^-/-^ UUO kidneys was significantly lower than that in WT kidneys from Day 3 (Day3: 32.9±2.9 vs.24.6±3.6 cell/field, P=0.026, Day5: 68.9±6.8 vs.45.1±7.4, P=0.049, Day7: 87.2±4.1 vs.66.3±0.3, P=0.002, WT vs. BLT1, respectively; Fig. 3B). Furthermore, expression of mRNA encoding F4/80 was significantly lower in BLT1^-/-^ UUO kidneys than in WT kidneys on Days 3, 5, and 7 (Day3: P<0.001, Day5: P=0.023, Day7: P=0.003, WT vs. BLT1, respectively; Fig. 3C).

**Figure 3.**
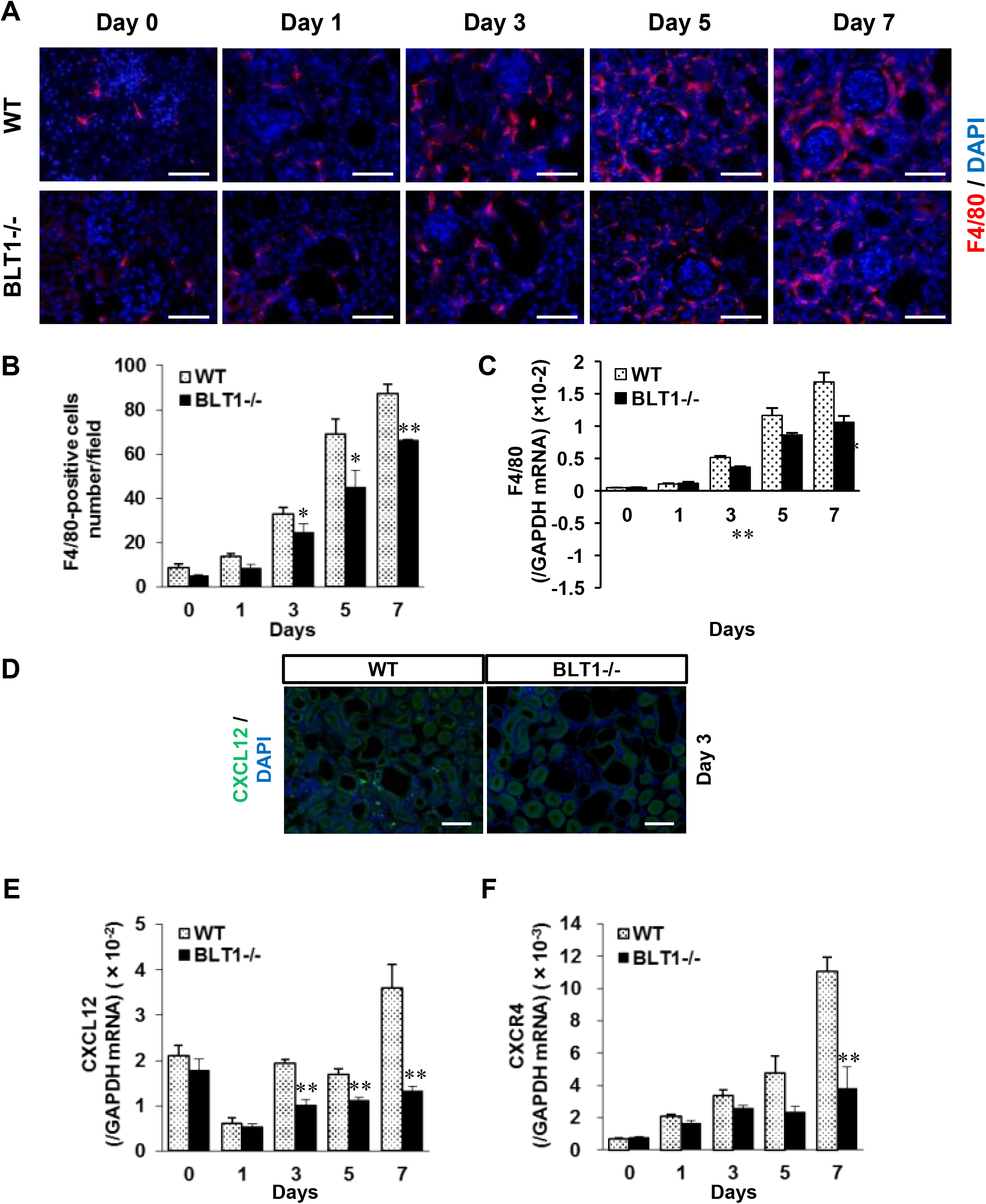
BLT1-induced accumulation of macrophages in UUO kidneys depends on the CXCL12/CXCR4 axis. A) Representative images showing F4/80 staining of WT and BLT1^-/-^ UUO kidneys. Scale bars = 50 μm. B) Changes in the number of F4/80-positive cells in WT and BLT1^-/-^ kidneys after UUO treatment. Data are expressed as the mean ± SEM (n=4 mice/group). C) Expression of mRNA encoding F4/80 in UUO kidneys from WT and BLT1^-/-^ mice. Data are expressed as the mean ± SEM (n=8-16 mice per group). *P<0.05 and **P<0.01, *vs*. WT mice. D) Representative images showing immunostaining of WT UUO kidney cortex with an anti-CXCL12 antibody (D) on Day 3. CXCL12-positive cells localized to the tubulointerstitial area. Scale bars = 200 μm. E,F) Expression of mRNA encoding CXCL12 (E) and CXCR4 (F) in UUO kidneys, as measured by real-time RT-PCR. Data are expressed as the mean ± SEM (n=8-12 mice per group). **P<0.01, vs. WT.

### Expression of CXCL12 in UUO kidneys is BLT1-dependent

Macrophages and fibroblasts are recruited to the kidneys following UUO [40]. Therefore, we examined chemokine levels in UUO kidneys from BLT1^-/-^ and WT mice. Immunofluorescence staining revealed that CXCL12 was expressed primarily in the interstitial spaces of WT UUO kidneys on Day 3 (Fig. 3D). Expression of CXCL12 mRNA in WT mice fell transiently on Day 1, before increasing again on Day 3; however, this increase was significantly lower in BLT1^-/-^ mice (Day 3,5,7: P<0.001, WT vs. BLT1, respectively; Fig. 3E). Moreover, expression of CXCR4, a specific ligand for CXCL12, in BLT1^-/-^ kidneys was significantly lower than that in WT kidneys on Day 7 (P=0.004, WT vs. BLT1, Fig. 3F). These results suggest that BLT1-induced macrophage infiltration into the UUO kidney is dependent on the CXCL12/CXCR4 axis.

### LTB_4_ increases expression of TGF-β, FGF-2, and collagen mRNA by macrophages *in vitro*

To evaluate whether expression of pro-fibrotic cytokines and collagen by macrophages is dependent on LTB_4_-BLT1 signaling, we isolated macrophages of WT mice from the peritoneal cavity and incubated them with LTB_4_. Expression of mRNA encoding TGF-β and FGF-2 increased 12 h after LTB_4_ treatment (Figs. 4A and B). Increased expression of collagen gene *Col1a1* was also detected 12 h after addition of LTB_4_ (Fig. 4C), although expression of αSMA did not increase after LTB_4_ treatment (Fig. 4D). These results suggest that LTB_4_ induces expression of collagen, TGF-β, and FGF-2 by macrophages.

**Figure 4.**
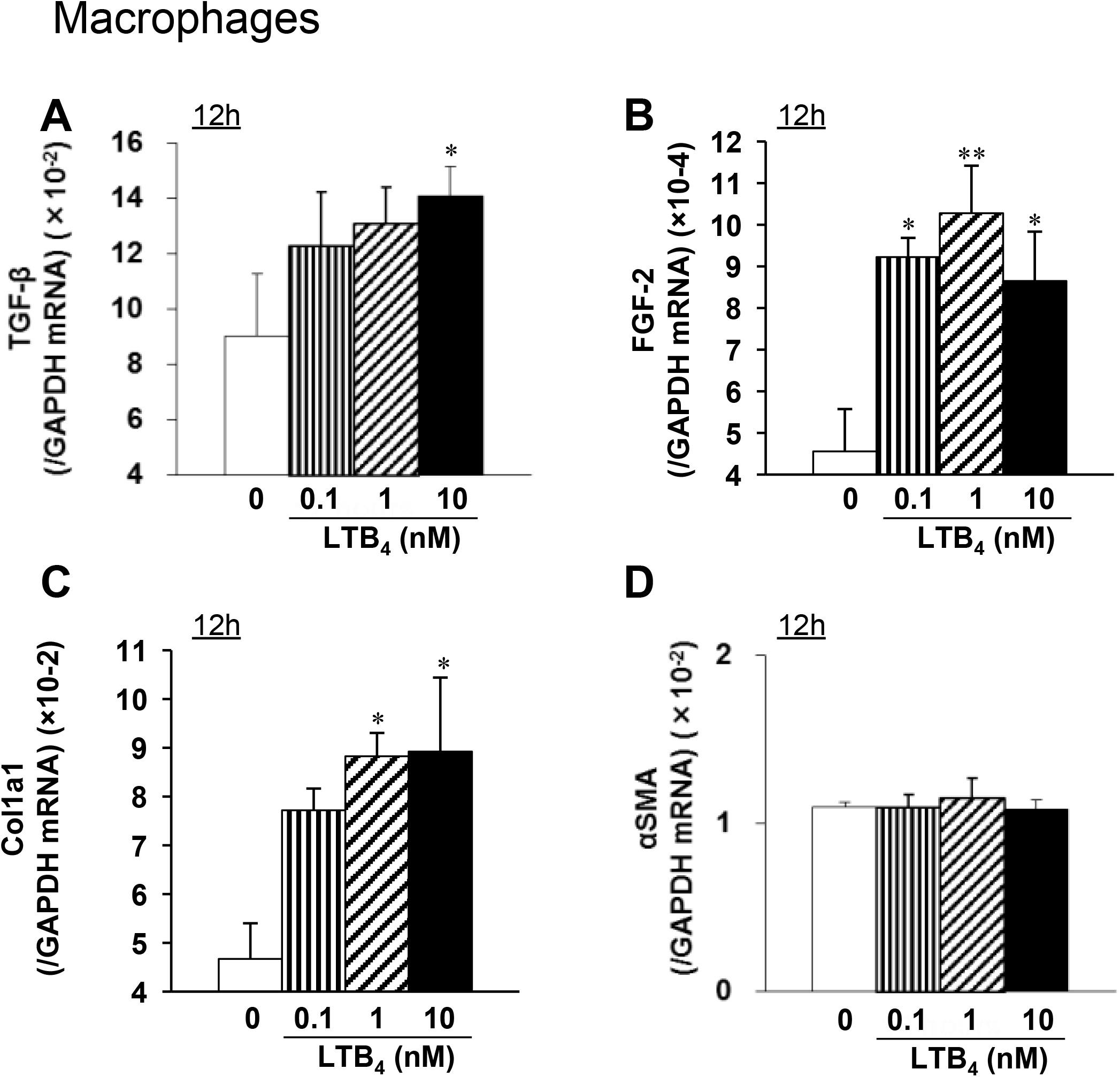
LTB_4_ increases expression of mRNA encoding TGF-β, FGF-2, and collagen by macrophages of WT mice *in vitro*. Macrophages collected from the peritoneal cavity of WT mice were incubated with LTB_4_ for 12 h. Expression of mRNA encoding TGF-β (A), FGF-2 (B), Col1a1 (C), and αSMA (D) was measured by real-time quantitative RT-PCR. Expression of mRNA encoding TGF-β, FGF-2, or Col1a1 (but not αSMA) increased after LTB_4_ treatment. Data are expressed as the mean ± SEM (n=6). *P<0.05 and **P<0.01, *vs*. 0 nM.

### LTB_4_ increases expression of mRNA encoding αSMA in connective tissue-derived mouse L929 fibroblast-like cells *in vitro*

When L929 murine fibroblasts were incubated with LTB_4_, we observed no increase in expression of mRNA encoding TGF-β or FGF-2 (Figs. 5A and B). Unlike in macrophages, LTB_4_ did not induce increased expression of mRNA encoding collagen 1a1, although αSMA mRNA levels increased slightly at 12 h after addition of 0.1 nM LTB_4_ (Figs. 5C and D). These results suggest that LTB_4_ does not trigger secretion of collagen 1, TGF-β, or FGF-2 by L929 cells. When L929 cells were stimulated with TGF-β, we saw no change in expression of mRNA encoding Col1a1 and αSMA (Figs. 5E and F).

**Figure 5.**
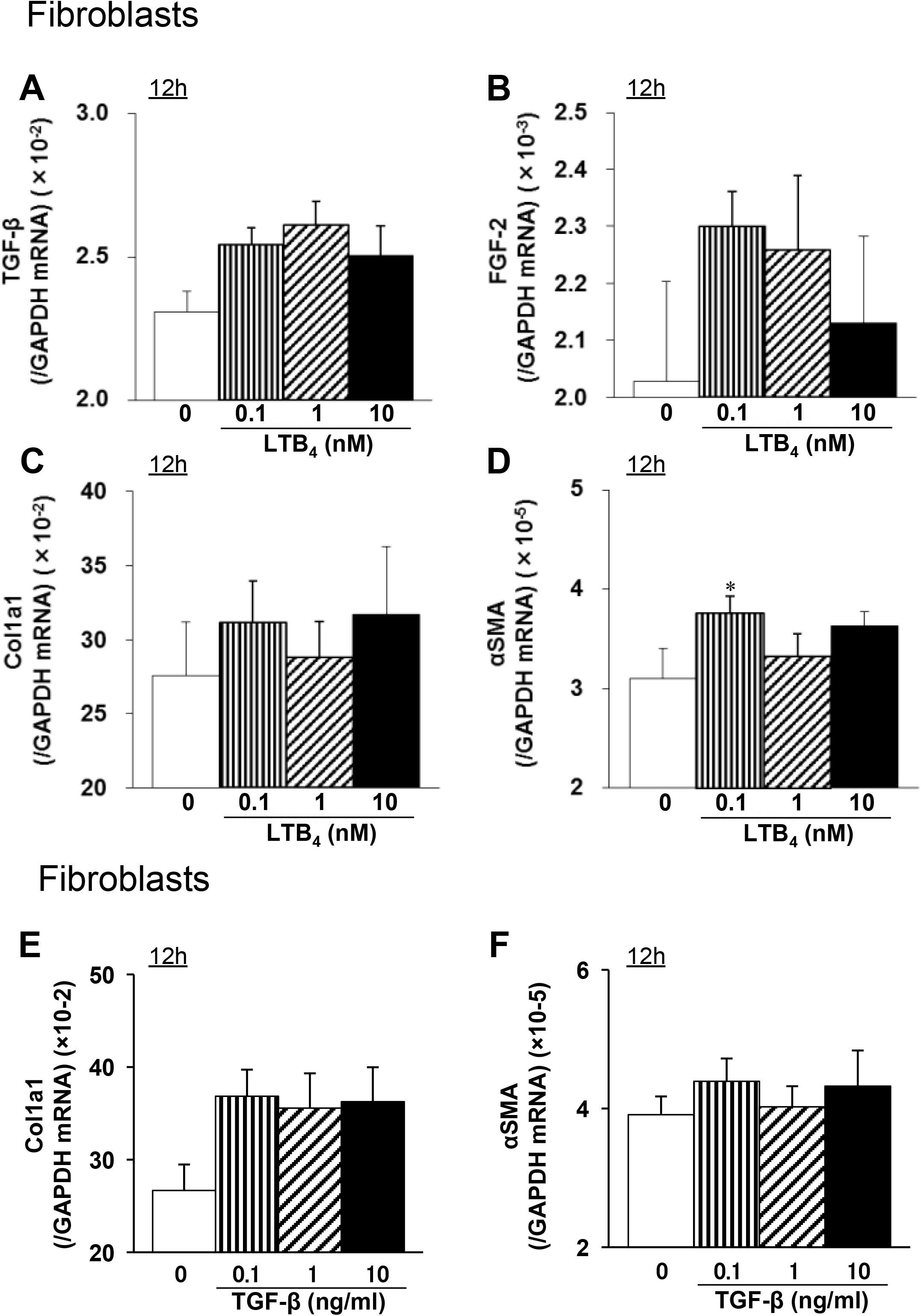
LTB_4_ increases expression of mRNA encoding αSMA in mouse L929 fibroblast-like cells derived from connective tissues. L929 mouse fibroblasts were treated with LTB_4_ for 12 h, and expression of mRNA encoding TGF-β (A), FGF-2 (B), Col1a1 (C), or αSMA (D) was measured by real-time quantitative RT-PCR. Data are expressed as the mean ± SEM (n=6). *P<0.05, *vs*. 0 nM. L929 mouse fibroblasts were treated with TGF-β for 12 h, and expression of mRNA encoding Col1a1 (E) and αSMA (F) was measured by real-time quantitative RT-PCR. Data are expressed as the mean ± SEM (n=6). *P<0.05, vs. 0 nM.

### BM-derived cells induced by LTB_4_-BLT1 signaling exacerbate fibrosis in the UUO kidney

Next, we examined whether BM cells induced by LTB_4_-BLT1 affect renal fibrosis. Selective deletion of the BLT1 receptor from the BM was performed by transplanting BM cells from BLT1^-/-^ mice. BM transplantation revealed that the area of type I collagen deposition in BLT1^-/-^-BM→WT UUO kidneys on Day 7 was significantly smaller than that in WT-BM→WT kidneys (14.5±0.5 vs.10.6±0.3 %, P=0.0006, WT-BM→WT vs. BLT1^-/-^-BM→WT, Figs. 6A and B). In addition, accumulation of S100A4-positive cells in BLT1^-/-^-BM→WT UUO kidneys at Day 7 was significantly lower than that in WT-BM→WT kidneys (52±4.7 vs.34.8±2.0 cells/field, P=0.015, WT-BM→WT vs. BLT1^-/-^-BM→WT, Figs. 6C and D). These results suggest that BM cells expressing BLT1 contribute to development of renal fibrosis in the UUO kidney.

**Figure 6.**
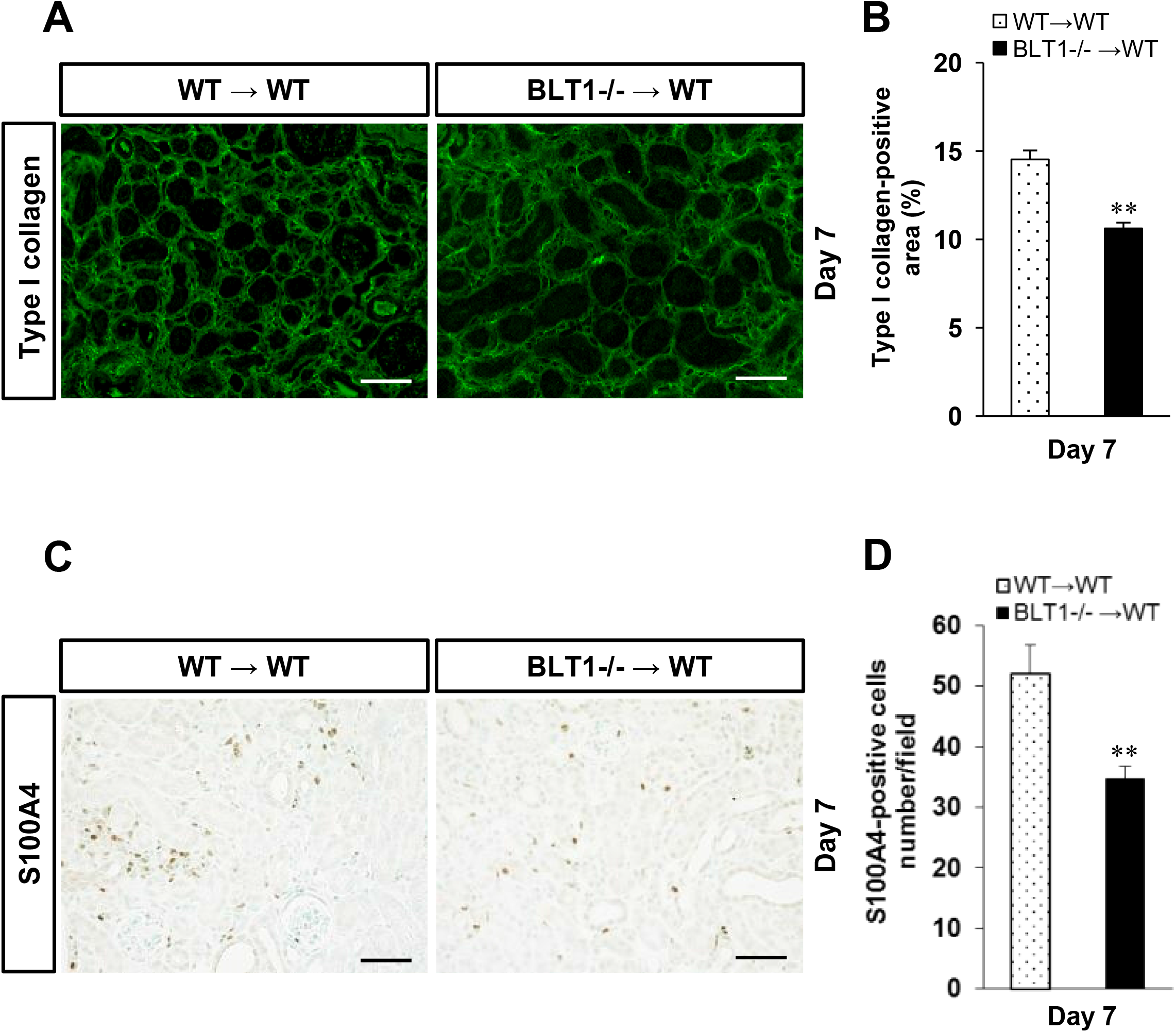
Bone marrow-derived cells induced by LTB_4_-BLT1 signaling exacerbate renal fibrosis in UUO kidneys. We examined renal fibrosis in UUO kidneys of WT mice transplanted with BLT1^-/-^-BM (BLT1^-/-^-BM→WT), and compared it with that in WT mice transplanted WT-BM (WT-BM→WT). A) Representative images showing type I collagen staining in the UUO kidneys of WT-BM→WT mice (left panel) and BLT1^-/-^-BM→WT mice (right panel) at Day 7. Scale bars = 200 μm. B) The type I collagen-positive area in UUO kidneys from WT-BM→WT mice and BLT1^-/-^-BM→WT mice at Day 7. Data are expressed as the mean ± SEM (n=4 mice/group). **P<0.01, *vs*. WT-BM→WT mice. C) Representative images showing S100A4 immunostaining in UUO kidneys of WT-BM→WT mice and BLT1^-/-^-BM→WT mice at Day 7. Scale bars = 200 μm. D) Number of S100A4-positive cells in WT-BM→WT mice and BLT1^-/-^-BMƒWT UUO kidneys. T Data are expressed as the mean ± SEM (n=4 mice/group). **P<0.01, *vs*. WT-BM→WT mice.

## Discussion

Here, we demonstrated that BLT1 signaling plays a major role in development of fibrosis in BLT1^-/-^ UUO model mice. Accumulation of collagen type I in UUO kidneys was significantly lower in BLT1^-/-^ mice than in WT. We also observed BLT1-dependent recruitment of macrophages and fibroblasts (Fig. 2 and Fig. 3); however, BLT1 signaling induced expression of mRNA encoding pro-fibrotic factors and collagen in a cell type-specific manner (Fig. 4 and Fig. 5). Surprisingly, we found that LTB_4_ acted on macrophages directly to upregulate expression of Col1a1. BLT1 signaling also induced expression of mRNA encoding TGF-β and FGF-2 *in vivo*, both of which are pro-fibrotic factors; however, we did not demonstrate production of TGF-β and FGF-2 by LTB_4_-stimulated fibroblasts *in vitro* (Fig. 5). BM transplantation experiments revealed that the area of type I collagen deposition in BLT1^-/-^-BM→WT UUO kidneys was lower than that in WT-BM→WT kidneys (Fig. 6). Thus, BM-derived cells induced by LTB_4_-BLT1 signals exacerbate renal fibrosis in the UUO kidney. Together, these results demonstrate a finely tuned mechanism underlying BLT1-dependent fibrosis in this model (Fig. 7), and suggest that blocking of BLT1 signaling may prevent fibrosis. The UUO model is good for studying tubulointerstitial fibrosis accompanied by cellular infiltration. LTB_4_ is a chemoattractant for leukocytes, particularly neutrophils and macrophages [21–24]. We found that, after UUO treatment, dilated tubular cells expressed 5-LOX (Fig. 1G), which may be important for initiating BLT1-dependent fibrosis. Expression of 5-LOX also induces LTB_4_, a chemoattractant for macrophages. These molecules elicit local extravasation of fibroblasts and macrophages, which infiltrate the tubulointerstitial space of the kidney in a BLT1-dependent manner.

**Figure 7.**
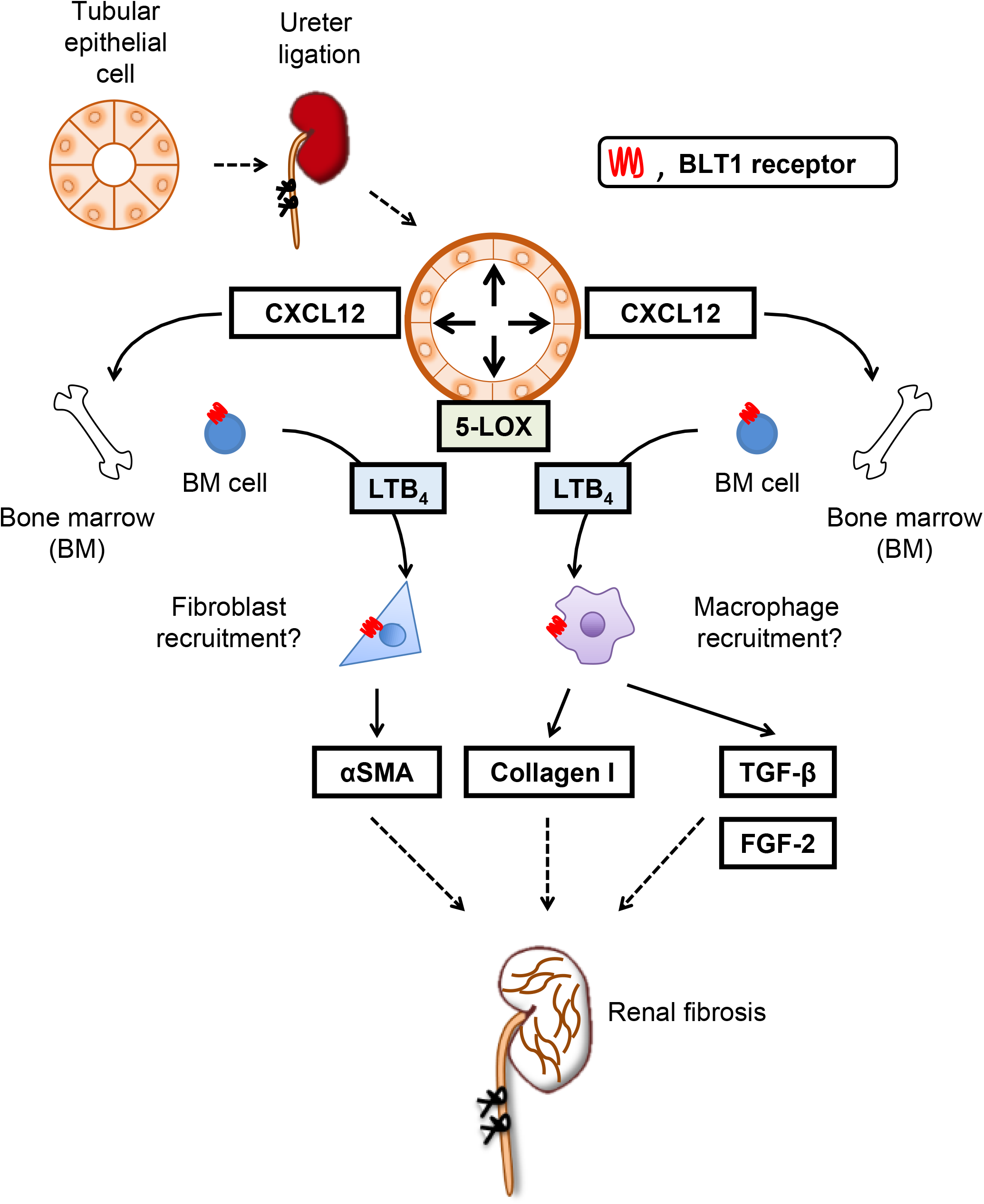
Role of LTB_4_-BLT1 signaling in UUO kidney fibrosis. BLT1 signaling is critical for development of fibrosis after UUO treatment. UUO treatment upregulates expression of CXCL12 in a BLT1-specific manner. LTB_4_ is also induced by UUO treatment. LTB_4_-BLT1 signaling increases recruitment of fibrocytes and macrophages to the kidney; these cells produce collagen, together with TGF-β and FGF-2, via BLT1 signaling, resulting in fibrotic changes.

PGE_2_ and PGI_2_, the most abundant eicosanoids, suppress fibrosis *in vivo* [18, 41]; however, few studies have examined involvement of LTs and other metabolites in fibrosis after UUO. Models of pulmonary fibrosis exhibit a synthetic imbalance favoring pro-fibrotic LTs over anti-fibrotic PGE_2_, suggesting a role for eicosanoids in fibrotic lung disease [42]. PGE_2_ levels in BAL fluid from patients with idiopathic pulmonary fibrosis (IPF) are lower than those from healthy control subjects [43]. In addition, fibroblasts grown from lung tissue isolated from patients with IPF synthesize less PGE_2_ than cells from healthy control subjects due to reduced COX-2 expression [44]. This has important pathologic consequences because decreased levels of COX-2 and PGE2 in these cells contribute to increased collagen synthesis and cell proliferation in response to TGF-β [44]. By contrast, BAL fluid from patients with IPF contains more LTB_4_ than that from control subjects [45]. LTB_4_ levels in lung tissue homogenates from patients with IPF are 15-fold higher, and those of LTC_4_ are 5-fold higher, than those in control subjects, reflecting constitutive activation of 5-LOX in alveolar macrophages [46]. Increased lung LT levels have been observed in mice after intratracheal administration of bleomycin; this is a commonly used animal model of pulmonary fibrosis [47]. In this model, fibrosis was blunted markedly following disruption of cPLA_2_ [48] and 5-LOX [47], suggesting that endogenous LTs play a major role in facilitating fibrosis. Several features of bleomycin-induced injury, including pulmonary recruitment of macrophages and neutrophils, alveolar septal thickening, fibroblast accumulation, and collagen deposition, are significantly less severe in LT receptor-deficient mice than in their WT littermates [49]. These results suggest that LTs are pro-fibrotic in these pathological settings. In contrast to lung fibrosis, there is no definitive evidence that LT is involved in UUO-induced fibrosis in the kidneys.

As mentioned above, we showed that 5-LOX expression increased after induction of UUO; we also observed 5-LOX-expressing cells in dilated renal tubules in UUO kidneys (Fig. 1G). A previous study in a mouse UUO model shows that COX-2 is upregulated in renal tubule epithelial cells after ligation of the ureter [18].

We observed BLT1-dependent accumulation of type I collagen, S100A4-positive fibroblasts, and αSMA-positive myofibroblasts in UUO kidneys (Fig. 2). Accumulation of S100A4-positive fibroblasts in the interstitial spaces within kidney tissues was evident from the early stages of UUO onset, although it was less marked in BLT1^-/-^ mice. The αSMA mRNA levels (Fig. 2E) suggest that αSMA-positive myofibroblasts accumulate in UUO kidneys; again, this was less marked in BLT1^-/-^ mice from Day 3. These results suggest that LTB_4_/BLT1 signaling is important for induction of renal fibrosis in UUO kidneys.

We also observed reduced accumulation of macrophages in BLT1^-/-^ mice during the early stage of UUO (Fig. 3). These accumulated macrophages supply TGF-β to sites of fibrosis. CXCL12 is a chemokine that plays a role in migration of BM-derived stem cells to the peripheral blood and from there to sites of tissue injury. CXCL12 is a potent chemoattractant for fibroblasts/myofibroblasts. CXCL12 binds to a specific receptor, CXCR4. Philips et al. report that the CXCL12/CXCR4 axis induces recruitment of BM-derived stem cells to injured lung tissue to induce pulmonary fibrosis [50]. Also, CXCR4 antagonists ameliorate renal fibrosis in the UUO kidney [51]. It would be worth investigating whether LTB_4_-BLT1 signaling interacts with the CXCL12/CXCR4 axis in the UUO kidney. Here, we found that CXCL12-positive cells localized primarily to the interstitial spaces within UUO kidneys (Fig. 3D). Furthermore, expression of CXCL12 and CXCR4 increased in WT kidneys more than in BLT1^-/-^ kidneys. These results suggested that the CXCL12/CXCR4 axis contributes to renal fibrosis in the UUO kidney in a LTB_4_-BLT1 signaling-dependent manner. The results also suggest that accumulation of collagen-producing cells is regulated by BLT1 signaling. Lack of BLT1 signaling may explain, at least in part, the reduced fibrosis observed in UUO kidneys. BM-mobilized macrophages and fibrotic cells express CXCR4 and infiltrate CXCL12-enriched tissue; however, we did not identify the type of cell that contributes to renal fibrosis in UUO kidney. Further experiments are needed to answer this question.

TGF-β has the potential to increase collagen biosynthesis by fibroblasts and myofibroblasts [52]. We confirmed reduced expression of TGF-β in UUO kidneys of BLT1^-/-^ mice, along with increased expression of TGF-β by macrophages stimulated with LTB_4_ *in vitro* (Fig. 4A). Interestingly, LTB_4_-stimulated L929 cells did not show increased expression of TGF-β and FGF-2 (Figs. 5A and B). Moreover, LTB_4_-stimulated macrophages upregulated expression of collagen 1a mRNA, but not that of αSMA mRNA (Figs. 4C and D). By contrast, LTB_4_-stimulated L929 cells upregulated expression of αSMA mRNA, but not collagen 1a mRNA (Figs. 5C and D). These results suggest that LTB_4_-BLT1 signaling induces fibrosis via accumulation of macrophages and fibroblasts. A previous report suggests that LTB_4_ does not alter expression of collagen-encoding genes in primary mouse lung fibroblasts. Furthermore, bleomycin-treated macrophages produce LTB_4_ and show increased production of TGF-β in a BLT1-dependent manner [53]. Taken together, these results suggest that LTB_4_-BLT1 signaling might promote TGF-β production by macrophages recruited via BLT1 signaling.

The results of the BM transplant experiments suggest that BM-derived cells induced by LTB_4_-BLT1 signaling exacerbate renal fibrosis in the UUO kidney (Fig. 6). Taken together, the results from the present experiments suggest that complex synergistic loops may be active during LTB_4_-induced fibrosis in this UUO model.

In conclusion, we show here that BLT1 signaling plays a role in development of fibrosis in UUO models. Accumulation of collagen type I in UUO kidneys of BLT1^-/-^ mice was significantly lower than that in UUO kidneys of WT mice. BLT1 signaling induced accumulation of macrophages and fibroblasts, and induced collagen biosynthesis directly via induction of TGF-β. Thus, BLT1-dependent fibrosis in this model is finely regulated, suggesting that BLT1 signaling is a good therapeutic target for preventing fibrosis.

## Acknowledgments

We thank Michiko Ogino, Naoko Ishigaki, Kyoko Yoshikawa, and Mieko Hamano for technical assistance.

BLT1: the high-affinity leukotriene B_4_ receptor
BLT1^-/-^: BLT1 knockout mice
UUO: unilateral ureteral obstruction
5-LOX: 5-lipoxygenase
CXCL12: C-X-C motif chemokine 12
CXCR4: C-X-C chemokine receptor type 4
TGF-β: transforming growth factor-β
FGF-2: fibroblast growth factor 2

